# Native Extracellular Matrix Orientation Determines Multipotent Vascular Stem Cell Proliferation in Response to Cyclic Uniaxial Tensile Strain and Simulated Stent Indentation

**DOI:** 10.1101/2020.12.11.421511

**Authors:** P.S. Mathieu, E. Fitzpatrick, M. Di Luca, P.A. Cahill, C. Lally

## Abstract

Cardiovascular disease is the leading cause of death worldwide, with multipotent vascular stem cells (MVSC) implicated in contributing to diseased vessels. MVSC are mechanosensitive cells which align perpendicular to cyclic uniaxial tensile strain. Within the blood vessel wall, collagen fibers constrain cells so that they are forced to align circumferentially, in the primary direction of tensile strain. In these experiments, MVSC were seeded onto the medial layer of decellularized porcine carotid arteries, then exposed to 10%, 1 Hz cyclic tensile strain for 10 days with the collagen fiber direction either parallel or perpendicular to the direction of strain. Cells aligned with the direction of the collagen fibers regardless of the orientation to strain. Cells aligned with the direction of strain showed an increased number of proliferative Ki67 positive cells, while those strained perpendicular to the direction of cell alignment showed no change in cell proliferation. A bioreactor system was designed to simulate the indentation of a single, wire stent strut. After 10 days of cyclic loading to 10% strain, MVSC showed regions of densely packed, highly proliferative cells. Therefore, MVSC may play a significant role in in-stent restenosis, and this proliferative response could potentially be controlled by controlling MVSC orientation relative to applied strain.

## 1 Introduction

Cardiovascular disease is the number one cause of death worldwide[1]. Stenting is the most common treatment for stenosed arteries, however 5-10% of all stents re-stenose[2]. In-stent restenosis is characterised by excessive cell proliferation, which may be driven by the proliferation of cells found in the media of an artery: vascular smooth muscle cells (VSMC)[3] and multipotent vascular stem cells (MVSC)[4]. VSMC have traditionally been thought to play a critical role in neointimal formation, whereby healthy, contractile VSMC transition to synthetic VSMC which proliferate and produce extracellular matrix within the intimal layer[5]. However, other more recent studies suggest that a small population of multipotent vascular stem cells (MVSC) migrate to the intimal layer and proliferate to form the neointima[4]. Notably, several studies have been conducted in an attempt to determine which cell population primarily contributes to neointimal formation, and while some studies demonstrated a VSMC origin[6; 7; 8], other studies have strongly indicated a MVSC origin for neointimal cells [4; 9; 10].

To-date, VSMC have been extensively studied and are known to be mechanically sensitive cells with the ability to change phenotype in response to mechanical forces, including mechanical strain[11; 12; 13; 14; 15]. In contrast, few studies have been done on MVSC. Recently, we have demonstrated that MVSC strain avoid and align perpendicular to the direction of strain in response to uniaxial tensile strain[16]. It must be noted, however, that although VSMC reorient when allowed to rotate freely on unstructured surfaces, in vivo VSMC are constrained by collagen fibers, where these fibers are oriented parallel to the dominant direction of strain[17; 18]. Therefore, while they cannot undergo strain avoidant reorientation in a healthy in vivo environment, they may have the opportunity to reorient in an environment where collagen fibers can remodel and change orientation, such as during vascular disease [19; 20]. For example, in mice, the development of atherosclerosis can cause collagen fibers to reorient to be less circumferential in direction[19]. Atherosclerotic plaques often have a paucity of collagen fibers within the lipid core, while fibrous plaque caps can show realignment of collagen fibers in the direction of the maximum principal strain[20]. Vascular disease interventions can also change the collagen fiber arrangement within a vessel; in an animal study on transapical valve replacement, stent strut contact with the pulmonary artery caused collagen fibers near the stent to align parallel to the stent struts, while leaving fibers between struts unaffected[21]. Additionally, in previous experiments from our lab, application of a simulated stent strut caused realignment of collagen fibers parallel to the stent strut direction[22]. These changes in collagen fiber orientation may also change the direction of vascular cells’ alignment, and consequently could influence the way in which these cells experience their strain environment. Despite MVSC being considered by many to play a critical role in vascular disease[23], to the authors’ knowledge, no prior study has investigated the role of native fiber structure on MVSC, and how fiber structure influences the response of MVSC to strain.

The aim of this study was therefore to investigate the response of MVSC to uniaxial tensile strain in the presence of native collagen structure, with fibers arranged either primarily in the direction of strain, or perpendicular to the direction of strain. To understand how the growth response to strain may indicate a role for MVSC in in-stent restenosis, we also designed and implemented a bioreactor system that simulated the loading of a single stent strut on strips of decellularized porcine carotid artery, recellularized with MVSC. This experimental setup enabled the response of MVSC to a stenting-type environment to be isolated from other vascular cell types and the contribution of MVSC to in-stent restenosis in a controlled strain environment to be more fully elucidated.

## 2 Materials and Methods

### 2.1 Cell Isolation and Culture

Rat MVSC were isolated by explant as previously described[16; 24] and were cultured in stem cell media containing high glucose DMEM with Glutamax (Bio-sciences), with 1% Penicillin/Streptomycin(Bio-Sciences), 1% N-2 supplement (Gibco), 2% B-27 supplement (Gibco), 20ng/mL bFGF (Corning), 2% chicken embryo extract (ATCC), 100 nM retinoic acid(Sigma-Aldrich), 50 nM 2-mercaptoethanol (Gibco). Rat MVSC were used at p19. All animal research was carried out in accordance with the principles of the EU Directive 2010/63/EU on the protection of animals used for scientific purposes through the Health Products Regulatory Authority (HPRA) of Ireland which upholds the principle of the “Three Rs” and makes it a firm legal requirement. All procedures were approved by the University Ethics Committee in accordance with the guidelines of the Health Products Regulatory Authority (HPRA) for the Care and Use of Laboratory Animals.

### 2.2 Decellularized Tissue

Porcine carotid arteries were dissected fresh and cryopreserved by freezing at a rate of −1°C per minute in tissue freezing medium (TFM) composed of 0.1M sucrose (Sigma-Aldrich) and 12.75% DMSO (VWR) in RPMI 1640 medium (Gibco). These arteries were then decellularized based on a previously determined protocol[25]. Briefly, they were perfused with 0.1 NaOH for 15 hours and then rinsed in saline for 65 hours. Vessels were then incubated in DNAse solution for 19 hours. Decellularized vessels were then returned to TFM and frozen until ready for use. Vessels were cut into 5mm x 10mm strips for static samples or 5mm x 15mm strips for strained samples, which had a strain region of length 10mm length. Strips were cut either circumferentially on the vessel for fibers predominantly parallel to strain direction, or axially on the vessel, for fibers predominantly perpendicular to strain direction,[26] as shown in Figure 1A. The intimal layer of the vessel was then removed using fine tipped forceps under a dissecting microscope to expose the underlying collagen fibres. Before cell seeding, strips were placed in 100% ethanol for at least 1 hour, then rinsed twice in sterile deionized water and twice in sterile PBS. Strips were then pinned flat as shown in Figure 1B and seeded with 1.33×10^4^ cells/cm^2^. Cells were allowed to adhere for 1 hour before strips were unpinned and transferred to culture for a further 3 days prior to initiating the application of strain.

**Figure 1:**
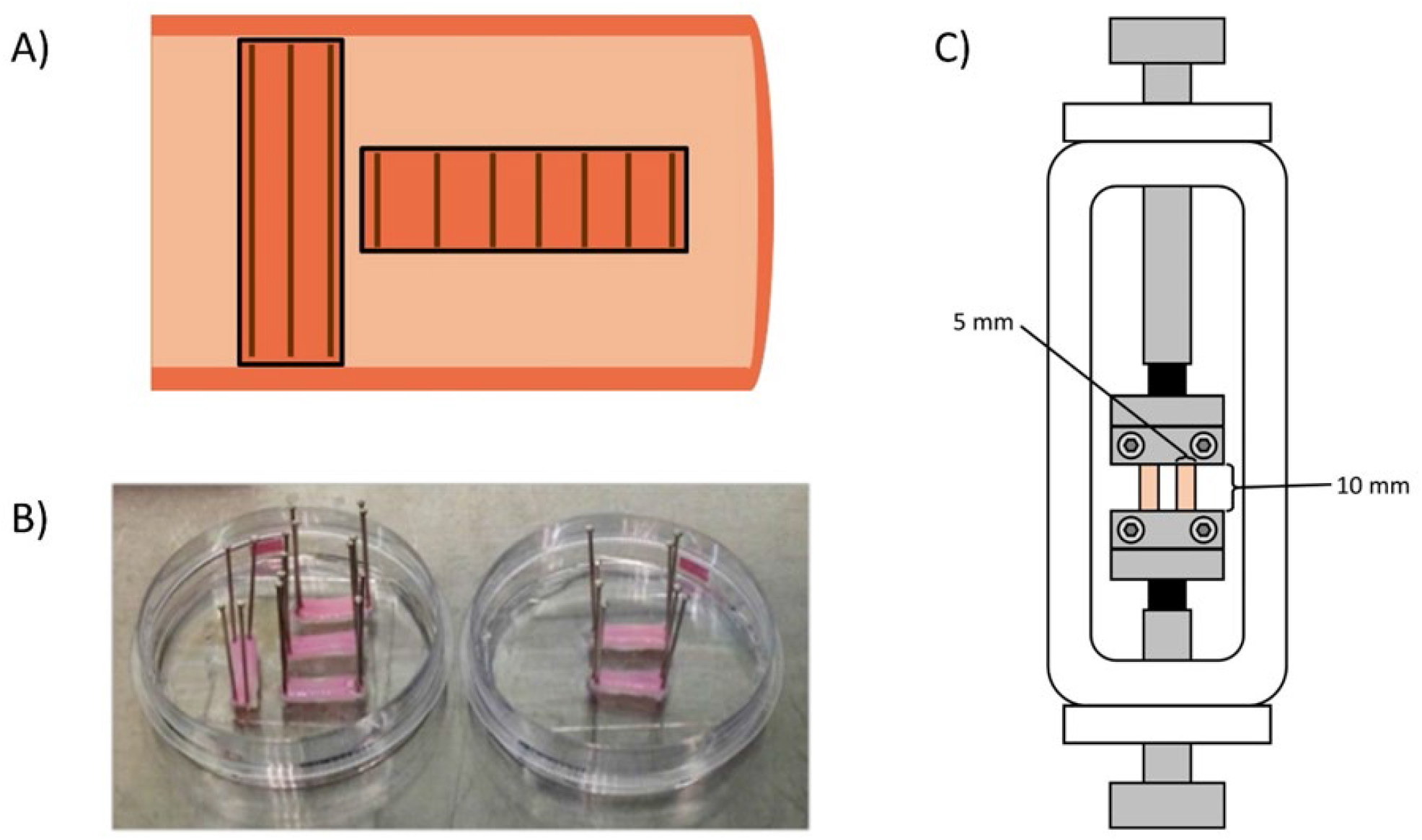
(A) Diagram showing how strips were cut from decellularized porcine carotid arteries in order to obtain strips with fibers predominantly oriented parallel or perpendicular to strain direction. (B) Strips pinned for cell seeding. (C) Setup of bioreactor chambers.

### 2.3 Application of Strain

For uniaxial tensile strain, strips were loaded into clamps within a Bose BioDynamic 5200 with two strips placed in each bioreactor chamber as shown in Figure 1C. Bose chambers were filled with high glucose DMEM with Glutamax (Bio-sciences) supplemented with 10% FBS and 2μl/mL Primocin. A preload of 0.05N was applied to all strips. Samples were strained uniaxially using displacement-controlled strain, from 0-10% strain at 1Hz for 10 days.

For simulated stent indentation, two indentation devices were melt-extrusion 3D printed in PLA. The static device shown in Figure 2 applied a static 10% elongation of the tissue, by indenting a 250μm wire 2.3mm into a recellularized tissue strip. The dynamic stenting device was designed to fit into a Bose Biodynamic 5000 system such that it could be indented to an arbitrary level. Strips were loaded into sterile rigs with two strips placed side by side as shown in Figure 3. Bose chambers or tubes were filled with 10% FBS medium supplemented with 2μl/mL Primocin. In the bioreactor, the wire indenter was moved so that it was just resting on the surface of the strips. The wire was then indented 2.3mm into the tissue strips at 1Hz for 10 days in order to apply a 10% strain to the tissue.

**Figure 2:**
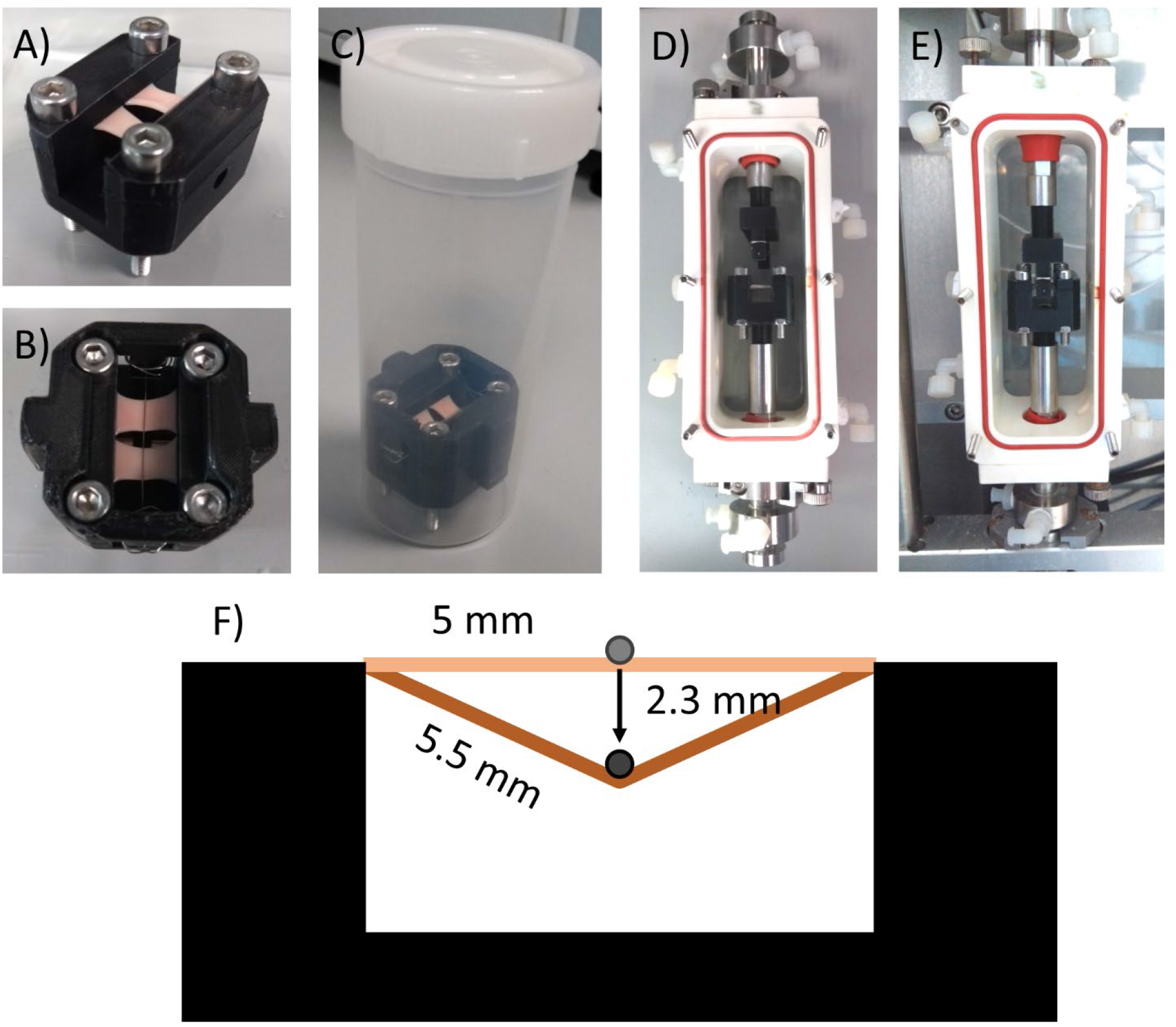
Images of static indentation device (A) Before application of stent strut, (B) Top view with stent strut, (C) In 120mL tube for addition of medium. Images of the dynamic indentation device (D) with strips loaded into the Bose bioreactor chamber, and (E) loaded into the Bose Biodynamic device with stent strut placed level with the top of the recellularized strips(F)Diagram showing 10% strain of tissue strips by stent strut application.

**Figure 3:**
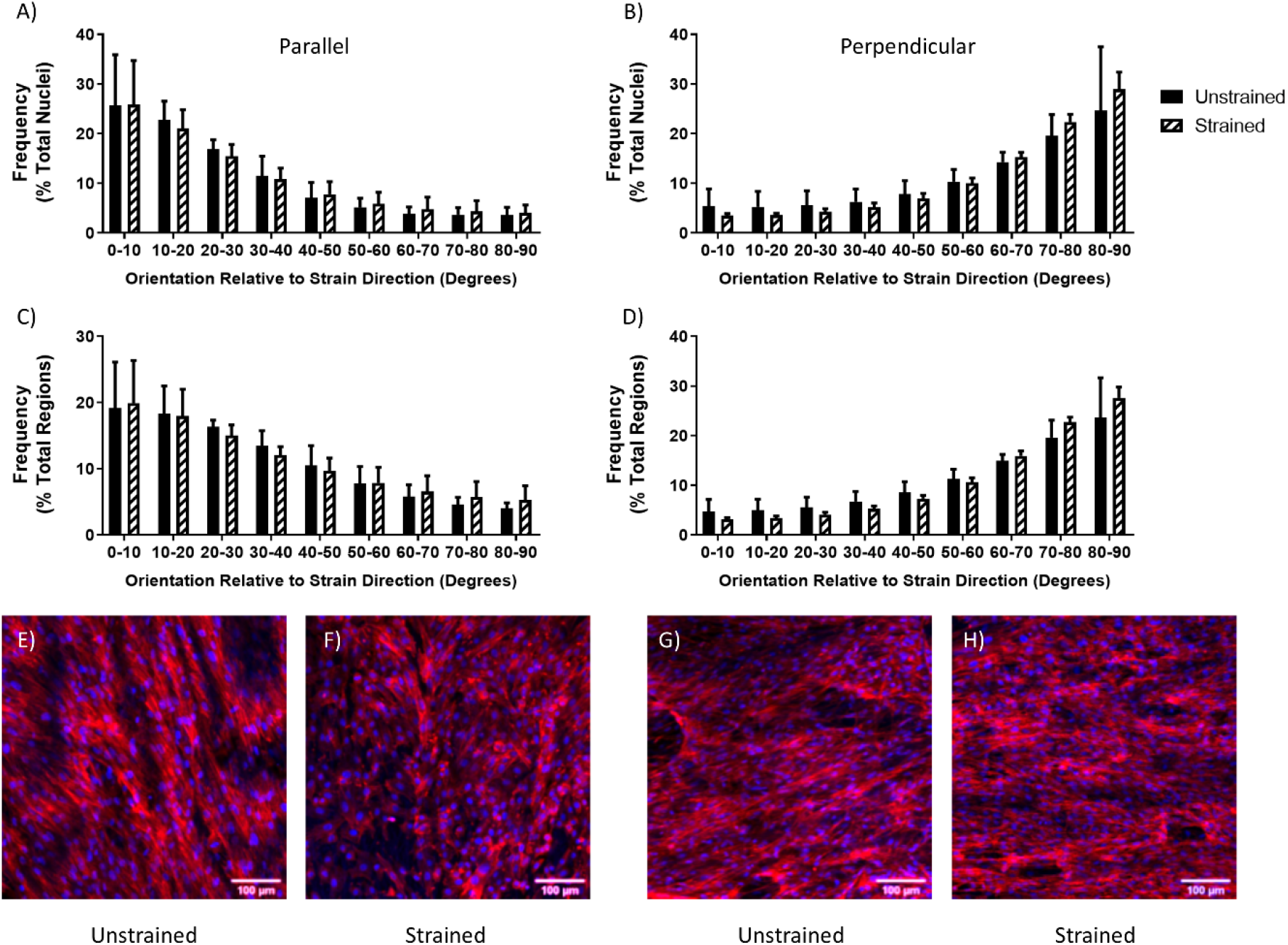
The alignment distribution of rMVSC (A,B) nuclei (C, D) and f-actin (C, D) on decellularized porcine carotid artery samples left unstrained, or (A, C) strained parallel or (B, D) perpendicular to the direction of collagen fibers. (E, F, G, H) Representative images of rMVSC on decellularized porcine carotid artery samples (E, G) left unstrained or (F) strained parallel or (H) perpendicular to the direction of collagen fibers.

### 2.4 Staining Protocol

Recellularized tissue strips were fixed for more than 24 hours at 4°C in 10% formalin. Strips were blocked and permeabilized in a solution of 5% BSA and 0.2% Triton-X 100 in PBS at room temperature for 40 minutes with agitation. Strips were then rinsed twice in PBS. Primary antibody solution was prepared (0.5% BSA, 0.2% Triton-X 100, PBS) and antibodies were used in accordance with the concentration information shown in Table 1. Strips were incubated in primary antibody overnight at 4°C with agitation. Strips were rinsed three times in PBS. Secondary antibody solution was prepared (0.5% BSA, 0.2% Triton-X 100, PBS) at a concentration of 1:1000, with 1:500 rhodamine Phalloidin, and 1:1000 DAPI. Strips were incubated in the secondary antibody solution in the dark for 48 hours at 4°C with agitation. Strips were rinsed three times in PBS and stored at 4°C protected from light before imaging. In indentation experiments, collagen was stained using collagen-binding adhesion protein 35 (CNA35)[27] prior to fixation. Strips were incubated in a 1:200 dilution of CNA35 in the medium used for strain application at 37°C for 2 hours, then rinsed twice in PBS before fixation.

**Table 1.**
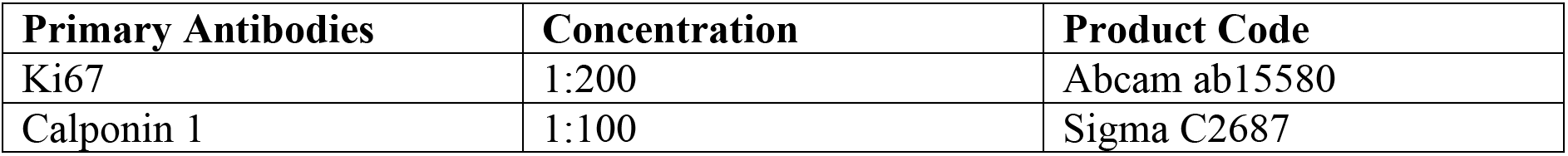

### 2.5 Imaging Protocol

All images were taken on a Leica SP8 scanning confocal microscope at 20x. For each sample, a continuous area of at least 24 images was scanned with a z stack taken at intervals of 10μm. The images were taken as 1024×1024 pixel images with a scan speed of 400 Hz.

### 2.6 Image analysis

Nuclear analysis was performed with ImageJ particle analysis to get nuclear number, nuclear orientation, major and minor axis, and nuclear area as previously described[16]. Images where nuclei were too dense to separate using particle analysis were counted and percentage of images that couldn’t be counted were calculated. Actin alignment was determined using the MatFiber Matlab program[28] which determined the alignment of fibers in each 5×5 pixel region of the image and enabled the % area of the tissue with a given degree of cell alignment to be ascertained. Ki67 nuclei were determined by masking Ki67 with nuclear area, and then counting positive nuclei using ImageJ particle analysis as previously described [16]. Images where Ki67+ nuclei were too dense to separate using particle analysis were counted and percentage of images that couldn’t be counted were calculated. Immunostaining intensity measures were determined as previously described[16], then normalized to a secondary antibody only control. Collagen fiber size range was determined by manually measuring the largest and smallest visibly distinct-CNA35 stained collagen fibers across multiple images of day 0 unstrained samples using the ImageJ ‘Measure’ function.

### 2.7 Statistics

Statistics were performed using GraphPad Prism 8.2.1. For experiments performed without simulated stenting, significance was determined using 2-way ANOVA with a Tukey’s post-hoc test or a Sidak’s post-hoc test as determined by the statistics software. When comparing across both parallel and perpendicular strain for both cell types a 3-way ANOVA was used with a Tukey’s post-hoc test. Each experiment had n ≥ 3 samples. For stenting experiments, significance was determined using an ordinary one-way ANOVA for cell counts and Ki67+ nuclear counts with n ≥ 3 samples. Collagen and F-actin alignment significance was determined using a 2-way ANOVA with a post-hoc Tukey test with n ≥ 3 samples.

## 3 Results

For rMVSC seeded on decellularized tissue strips, application of 10%, 1Hz uniaxial tensile strain for ten days resulted in cells aligned with the direction of collagen fibers regardless of whether the cells were oriented parallel or perpendicular to the direction of strain. This cellular orientation was determined both by the direction of the major axis of the nuclear ellipse and by the orientation of f-actin cytoskeletal fibers (Figure 4). In order to determine how strain influenced the overall number of cells, cell nuclei were counted to determine cell number. There was no significant change in cell number between cells that were left unstrained, or cells that were strained either parallel or perpendicular to fiber direction (Figure 5A). Cells were also assessed for proliferation by counting the number of Ki67 positive nuclei, which would indicate an actively dividing cell. Cells which were exposed to strain parallel to fiber direction showed an increased percentage of Ki67 positive nuclei compared to cells that were strained perpendicular to fiber direction (Figure 5B).

**Figure 4:**
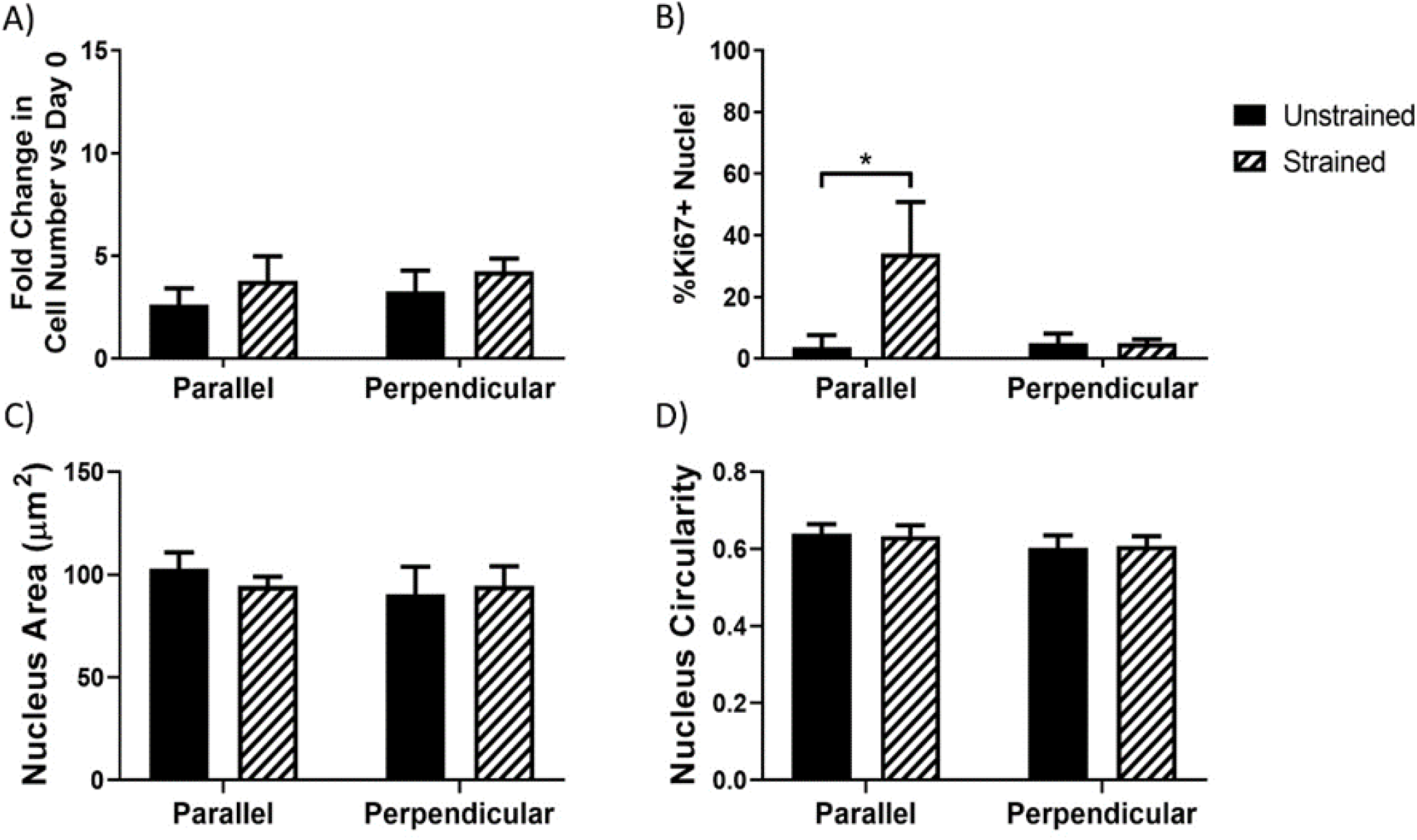
(A) Fold change in cell number, (B) percentage of Ki67 positive cells, (C) nuclear area, and (D) circularity of the nucleus, measured by the short axis divided by the long axis of the nuclear ellipse for both rMVSC strained parallel or perpendicular to collagen alignment direction. * p < 0.05

**Figure 5:**
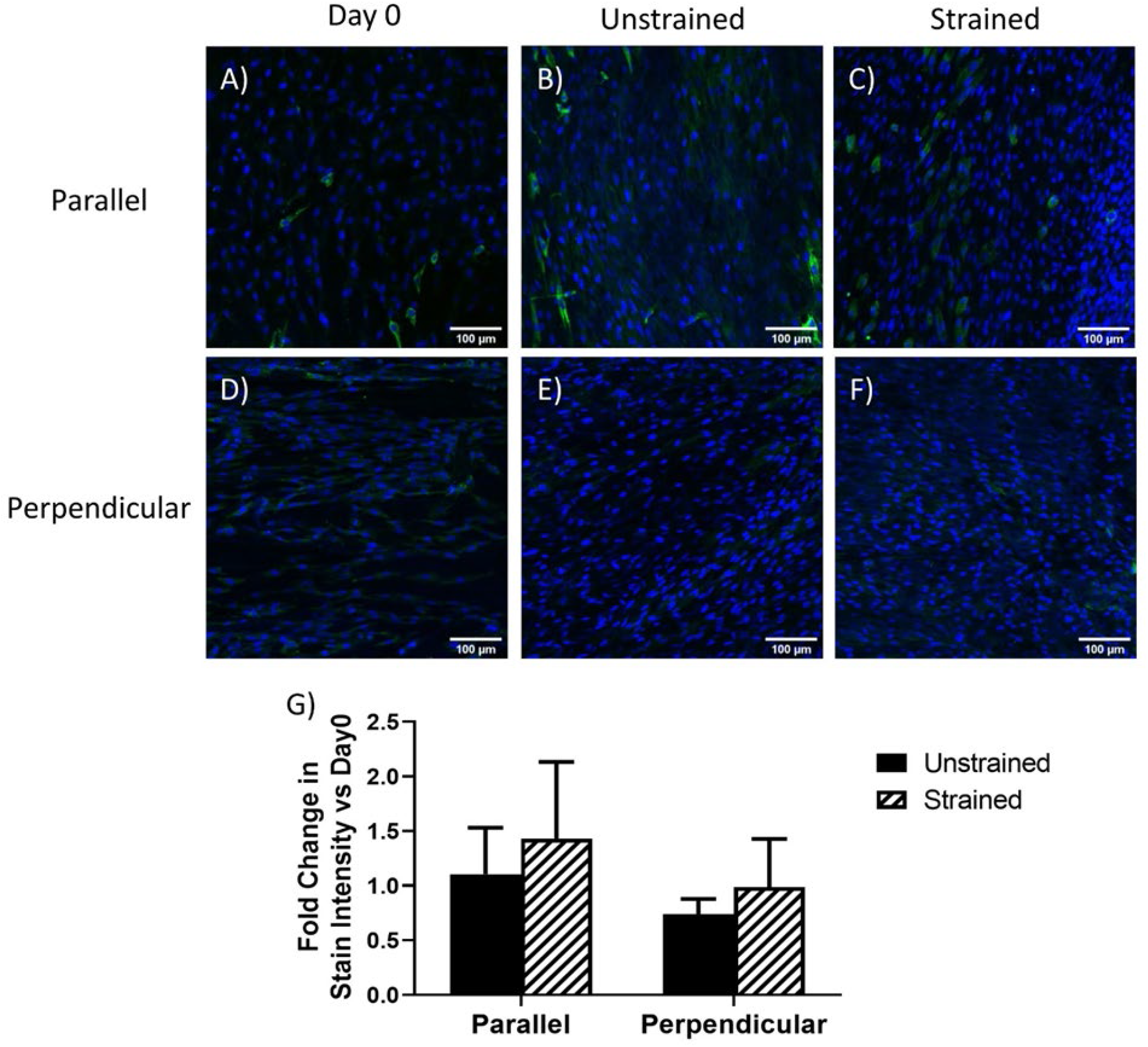
Representative images of rMVSC stained for nuclei (blue) and Calponin 1 (green) (A, D) before strain, (B, E) after 10 days without strain, or 10 days of 10%, 1Hz uniaxial tensile strain (C) parallel or (F) perpendicular to fiber direction. (G) Stain intensity analysis showed no significant differences between conditions.

Cell nuclei were also assessed on their size and shape. Cells showed no significant changes in nuclear size (Figure 5C) or nucleus circularity (Figure 5D) in response to strain. Cells were assessed for calponin 1 protein expression in order to determine if either cell was adopting a contractile VSMC phenotype. None of the cell types showed a significant change in calponin 1 expression in response to strain either perpendicular or parallel to fiber direction (Figure 6).

**Figure 6:**
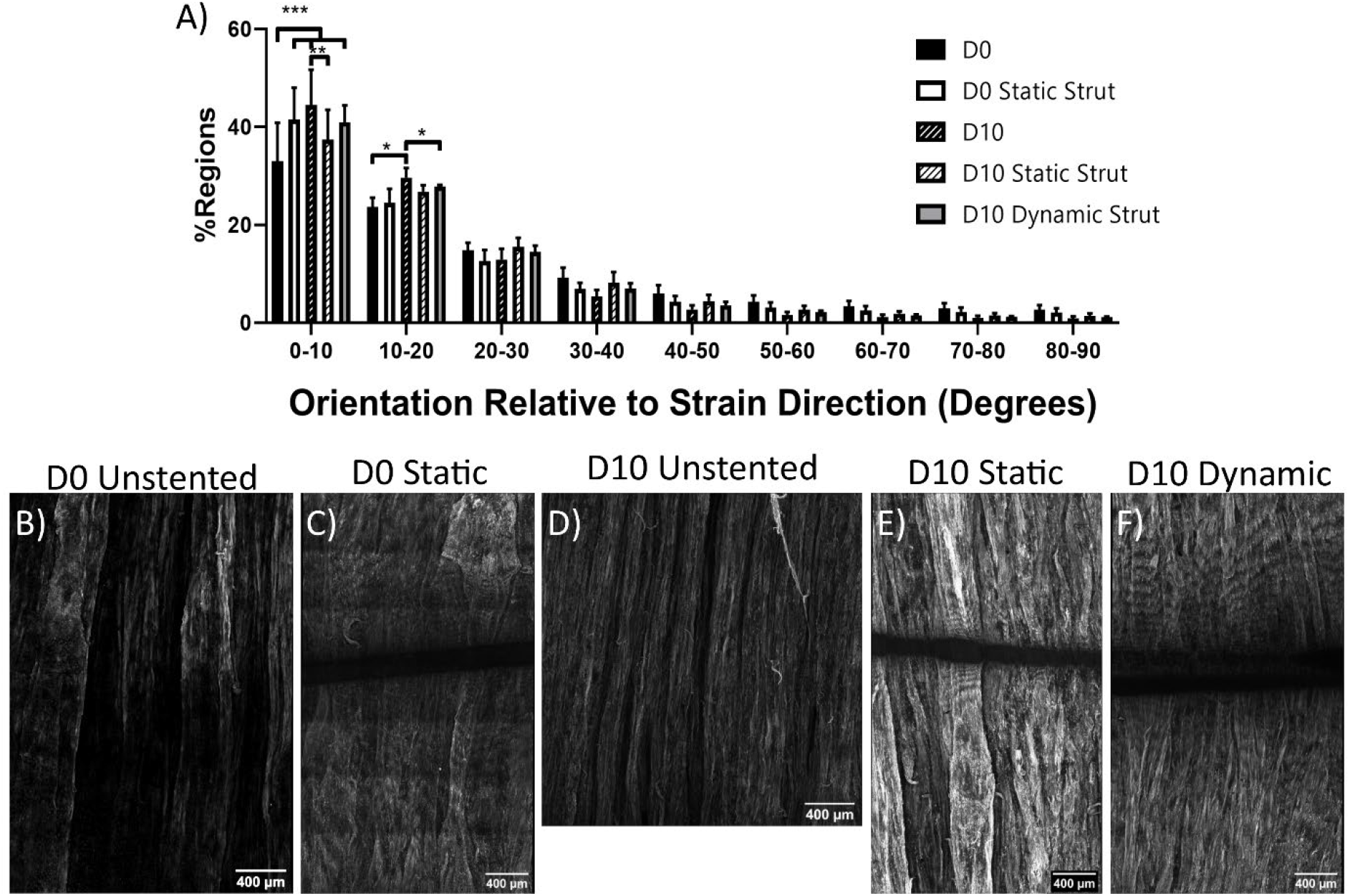
Orientation of collagen structure of decellularized porcine carotid artery relative to strain direction. (A) Fiber distribution changes after stent strut loading regimes. (B-F) Representative images of collagen fibers stained with CNA35. Grey bars in C, E and F indicate stent location. Scale bars = 400μm *p<0.05, **p<0.01, ***p<0.001, ****p<0.0001

In the stent indentation model, circumferentially cut strips were used so that collagen was aligned primarily parallel to the direction of strain and collagen was stained in order to determine if stent strut loading would cause collagen reorientation. Overall, all conditions showed collagen aligned primarily parallel to the direction of strain (Figure 7). Collagen fiber size in the day 0 unstrained samples ranged between 0.5 to 4μm in diameter. After ten days, cell alignment was evaluated using f-actin staining of cell actin cytoskeleton. Ten days of loading with either a static or a dynamic stent strut decreased the alignment of cells parallel to the direction of the collagen fibers and strain (Figure 8). In samples without a stent strut, cells remained aligned with the direction of collagen fibers (Figure 8A). In both statically loaded and dynamically loaded samples, cells aligned along the stent strut, perpendicular to the direction of the collagen fibers (Figure 8B, C). Further away from the stent, however, in statically loaded samples cells remained aligned with the underlying collagen fibers (Figure 8D), while in dynamically loaded samples, some regions showed cells aligning perpendicular to the direction of strain and collagen fibers (Figure 8E).

**Figure 7:**
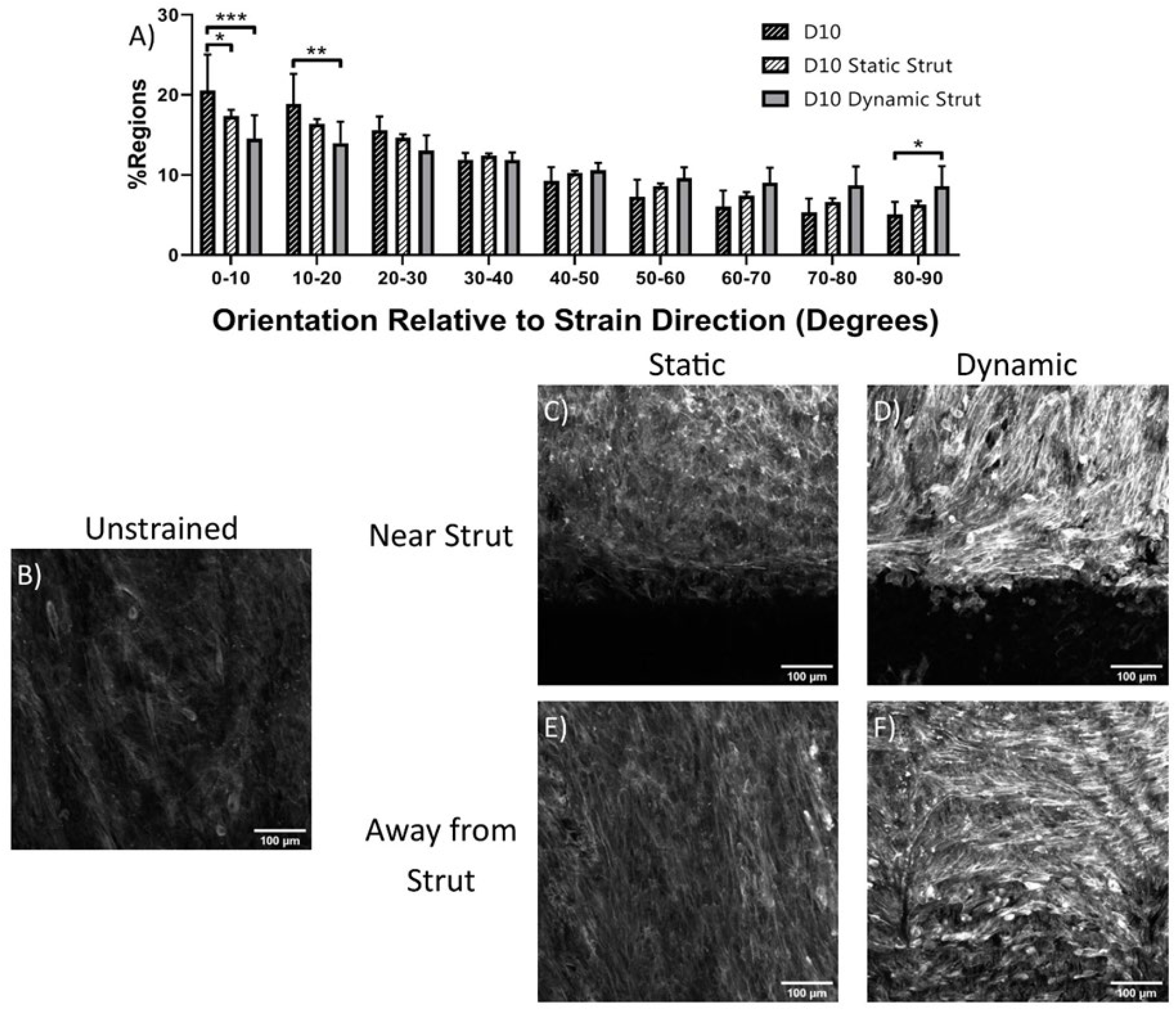
F-actin orientation of cells exposed to various loading regimes for 10 days. (A) Stent strut indentation decreases cell alignment in strain direction. Representative images of cells stained for f-actin for (B) unstrained and (C, E) statically and (D, F) dynamically loaded cells, both (C, D) near and (E, F) far from stent strut. Scale bar = 100 μm *p<0.05, **p<0.01, ***p<0.001, ****p<0.0001

**Figure 8:**
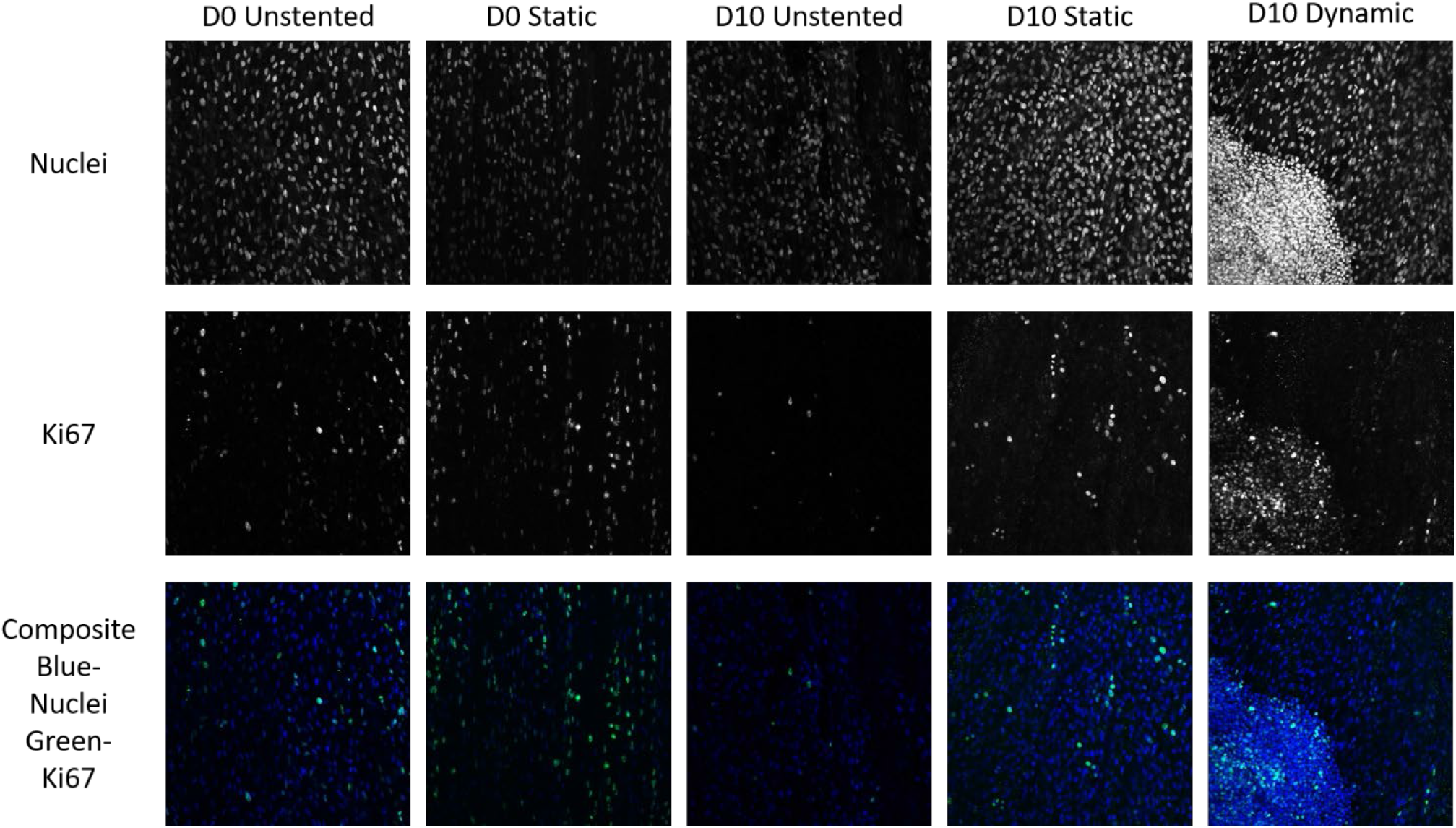
Representative images of nuclei and Ki67+ nuclei for different stent strut loading regimes. Scale bar = 100 μm

Cell proliferation was evaluated both by cell count and by the number of Ki67 positive cells. While samples from day 0, and unstrained cells at day 10 had countable numbers of nuclei, and Ki67+ cells (Figure 9A-C,F-H,K-M), samples that were exposed to a static stent strut for 10 days had many frames of cell nuclei that were too dense to count (Figure 9D). Samples that were exposed to a dynamically loaded stent strut for 10 days had both regions of nuclei, and regions of Ki67+ nuclei that were too dense to count (Figure 9E, J, O).

**Figure 9:**
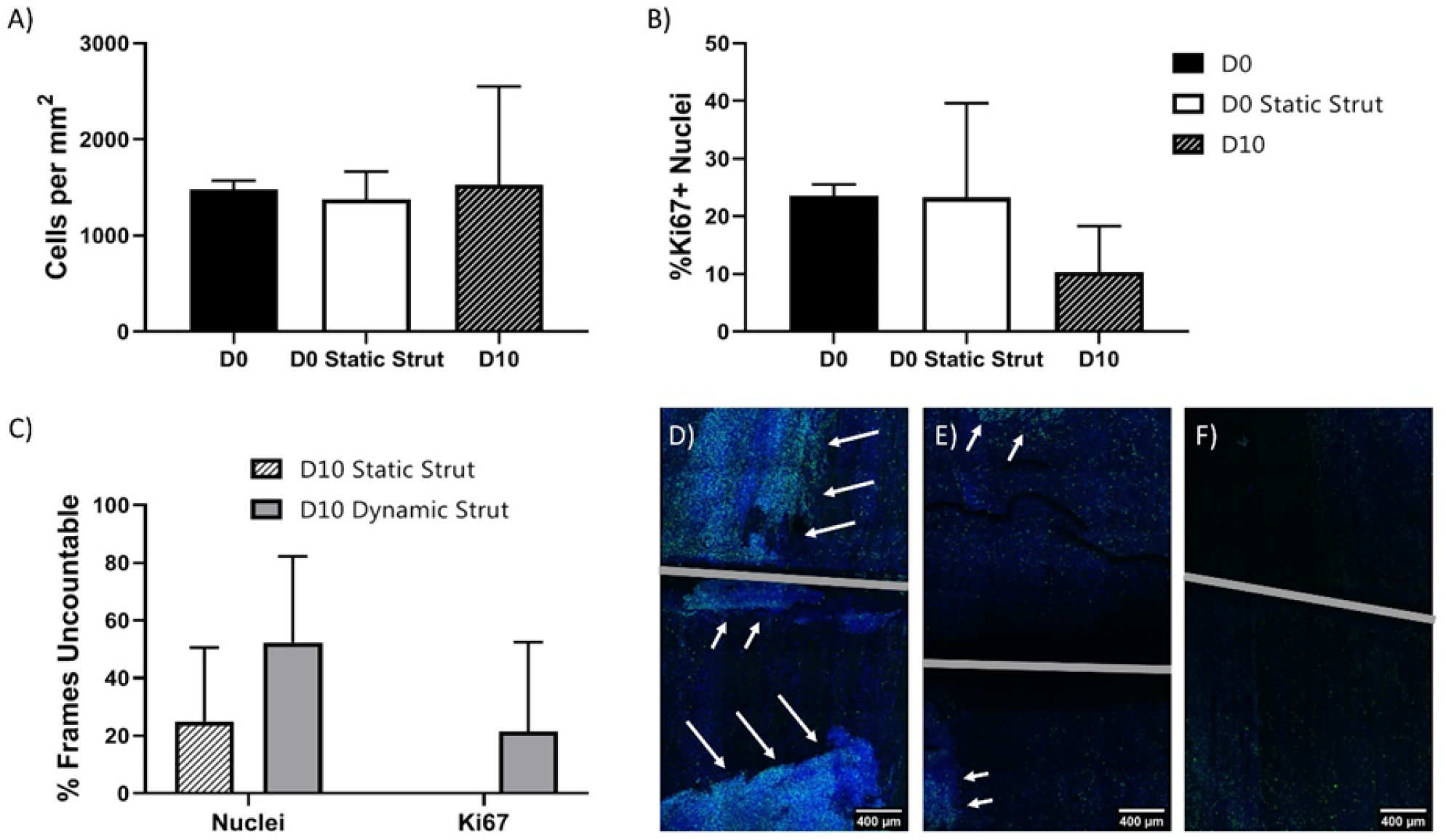
Cell number and proliferation. (A) Cell number for conditions with countable nuclei. (B) Percentage of Ki67+ nuclei for conditions with countable nuclei. (C) For conditions without countable nuclei, percentage of frames for which individual nuclei or Ki67+ nuclei cannot be counted. (D-F) Images of dynamically loaded samples, with arrows pointing to regions of high density, highly proliferative cells. Scale bars = 400μm

When the nuclear staining results were quantified, neither cell number nor percentage of Ki67+ nuclei was significantly different between either of the day 0 conditions or day 10 samples that were left unstrained (Figure 10A, B). Both day 10 conditions that involved loading with a simulated stent strut had regions where cells were too densely packed to count nuclei. The number of uncountable frames was slightly, but not significantly higher in the dynamically loaded samples than the statically loaded samples, however only dynamically loaded samples had regions where Ki67+ cells were too densely packed to count (Figure 10C). Most of the samples that were dynamically loaded had regions of densely packed and highly proliferative cells (Figure 10 D, E, F).

## 4 Discussion

This paper is the first to investigate the effects of both structure and tensile strain on MVSC, a cell type that has been shown to play a role in intimal hyperplasia. In stenting situations, however, tensile strain is much more complex than simple uniaxial loading. Therefore, a custom-designed test rig was developed to simulate the indentation of a single stent strut. This device allowed for the generation of strains in vascular tissue similar to those found in an in vivo stenting environment.

### 4.1 Alignment

The aligned structure of the medial layer of the decellularized artery induced both nuclear and cytoskeletal alignment in the direction of fiber alignment in MVSC. While few studies on MVSC microstructural response have been performed, MVSC were shown to align in the direction of PDMS microgrooves[29]. VSMC microstructural response may also give some indication of how MVSC would respond to microstructural cues. VSMC have previously been shown to align parallel to structural cues including microgrooved collagen substrates,[30; 31] micropatterned PDMS, [32; 33] and aligned electrospun[34] and melt-spun fibers.[35] The native collagen structure causes cell alignment in a similar manner to engineered patterned surfaces despite having a less controlled topography. The collagen fibers in our decellularized substrate act in a similar manner to these synthetically patterned substrates. Therefore, these decellularized vessels can be used to prime vascular cells to align in a circumferential direction, just like the VSMC in a native artery.

When exposed to strain, MVSC remained constrained by structure. The lack of strain avoidant reorientation shows that structural cues will dominate over strain cues. Because structural cues can prevent strain-induced orientation changes, tissue engineering applications or other applications that change collagen alignment can use microstructural cues to determine MVSC alignment regardless of applied strain. However, the ability of structure to influence cell response to strain may depend on the spacing of the cues provided. One study on VSMC indicated that microgroove size could determine the extent that microstructural cues dominate over strain cues. VSMC on both the thickest 70 μm microgrooves and the thinnest 15 μm microgrooves showed more strain avoidant realignment than in the middle microgroove width of 40 μm[32]. This indicates that there is a sweet spot of microgroove width in which cells will remain the most aligned, even when subjected to uniaxial tensile strain.

Loading recellularized strips with a simulated stent strut also changes cell alignment. Although previous experiments showed some collagen reorientation in response to the loading of stent struts,[21; 22] in this simulated stent loading regime, no detectable collagen reorientation occurred. However, in previous experiments this reorientation occurred below the surface level of the vessel,[22] whereas this experiment only observed the inner-most surface layer of collagen. As a result, it may be that the collagen reorientation is occurring at a deeper level where it would not affect the cells that were seeded only on the surface of the vessel strips.

Even without collagen reorientation, cell re-alignment was observed in samples that were loaded by the simulated stent strut for 10 days. The cells that realigned along the length of the stent strut were likely aligned due to contact guidance from the edge of the stent strut. Because *in vivo* stenting causes de-endothelialization, any cells that migrate to the intimal space would contact the stent strut itself, changing the cellular orientation. This change in orientation could change how the cells experience strain. Under dynamic stent strut indentation, some regions of cells not near the stent strut realigned perpendicular to strain direction. This is not unexpected, given that MVSC that aren’t constrained by underlying microstructural cues will reorient perpendicular to strain direction[16]. In regions where cells reoriented perpendicular to fiber and strain direction, the cells that remained in contact with the ECM structure remained oriented with the fiber direction, while the cells that had reoriented were layered over the top of other cells, so they were no longer sensing the microstructural cues from the underlying ECM. If MVSC are the primary driver of in-stent restenosis, based on the results presented here, it would be anticipated that the cells proliferating in in-stent restenosis would be in regions where there is ECM or other guidance causing cells to orient parallel to the direction of strain.

### 4.2 Proliferation/Apoptosis

When exposed to strain parallel to the direction of cell alignment, the number of dividing, Ki67+ MVSC increased. MVSC on unstructured substrates have shown a decrease in cell number in response to 10%, 1 Hz uniaxial cyclic tensile strain[16]. However, in these experiments, cells were free to reorient and strain avoid. When an elastic cell substrate is strained parallel to cell alignment, the cell itself deforms to a greater degree[36]. Cell alignment can be critical to obtaining the desired cell response. When MVSC were strained parallel to cell orientation, cell proliferation increased. Interestingly, these strain-induced changes did not appear when the cells were aligned perpendicular to the direction of strain, showing that cell orientation relative to strain direction is critical for how the cells sense uniaxial tensile strain. It is also worth noting that whilst cells aligned in the direction of strain increased in proliferation, they did not significantly increase in cell number. Because cell proliferation increased while cell number did not significantly increase, it is likely that MVSC death is also increasing when the cells sense strain. A significant finding in this work is that cells only appear to sense the strain when they are aligned with the direction of the strain.

When samples were loaded with a stent strut for 10 days, both the static strut and the dynamic stent induced an increase in cell number versus unstrained samples. While static strain has not been investigated in MVSC specifically, static strain in intact tissue rings showed increased DNA synthesis, possibly indicating increased cell division, in medial vascular cells [37]. This could indicate the presence of MVSC in these intact tissue samples or indicate that VSMC respond differently to static strain than cyclic tensile strain. Previous studies have shown differing responses for VSMC between static and cyclic strain. VSMC under stationary strain increased MMP-2 and MMP-9 expression, while cyclic strain decreased production of these ECM degrading proteins [38]. In this experiment, we see that whilst static strain increased cell proliferation, it is still qualitatively different than the proliferation seen in cyclically loaded samples.

Given the increased proliferation of MVSC when strained in the direction of cell alignment, we would have expected to see an increase in MVSC number when they were dynamically loaded parallel to the strain direction. When dynamically loaded, samples began to show regions of dense, highly proliferative cells. These areas mimic areas of in-stent restenosis which have both high cell density [39], and high proliferation [40]. This study correlates with studies of various stent designs that indicate that designs that cause increased strains within the vessel cause increased in-stent restenosis [41; 42; 43]. Additionally, many parameters that might induce greater strains within the vessel have been shown to increase the extent of in-stent restenosis including more rigid stents [44], and stents that are mismatched to the size of the artery[45]. The dynamic strut indentation simulates a loading environment where there is high dynamic strain applied within the artery, which would be consistent with the regions that showed high in-stent restenosis.

While only the dynamic stent indentation showed the development of dense, highly proliferative cell regions, static stenting also showed an increase in overall cell number compared to unstrained samples. Therefore, both types of loading might increase neointimal thickness to some degree. It is possible that the differences in how the cells perceive static versus cyclic strain are due to how the different types of strain affect mechanotransduction through the actin-cytoskeleton. In MSC, cyclic strain causes a greater rearrangement of the cytoskeleton of cells stretched statically to the same strain [46]. These differences in static versus dynamic response have implications in stent design. Stents that allow the vessel to drape, or prolapse, within the stent may be at higher risk of developing in-stent restenosis than stents that hold the blood vessel taut, if MVSC are the key cell type in neointimal formation. Stent tissue prolapse has previously been shown to be higher in bare metal stents with higher restenosis rates[47].

### 4.3 Applications

This experiment also shows how critical cell alignment is in tissue engineering applications. In order to induce a specific behavior in cells in response to tensile strain, the cells must first sense the strain. Therefore, knowing the orientation of the cells within the tissue engineered construct is important in selecting the correct strain amplitude to apply to achieve the desired phenotypic and proliferative results. The combination of MVSC sensitivity to strain amplitude previously described [16], and strain orientation relative to cell orientation means that varying these two parameters can be used in order to direct vascular cell and tissue development into the desired structures. Multi-scale mechanical modelling could then be used to predict how the mechanical properties of the vascular tissue change as the cells grow and differentiate in order to design a loading regime that would be able to adapt as the tissue matures.

While this research makes a strong argument for MVSC being a key player in in-stent restenosis, the device that was designed to provide stent strut indentation could be used to investigate many further parameters of stent design. Since strut thickness has been shown to influence the restenosis rate[48], stent strut diameter and cross-sectional shape could be modified by swapping different wires into the indentation devices. In addition, stent struts are often oriented in directions other than axial to the vessel, and we clearly demonstrated that ECM and cell alignment relative to strain direction plays a vital role in how cells sense strain. By cutting decellularized tissue strips at angles other than circumferential to the artery, the effect of stent strut orientation could be studied allowing for a better understanding of how to control stent strut orientation in order to prevent in-stent restenosis. With some minor modifications to the static device, and only modifying input parameters on the dynamic device, indentation depth could be altered in order to apply different levels of strain to the strips. This would clearly be of interest given that previously we have demonstrated that different amplitudes of uniaxial strain can influence cell response to strain[16]. This information could be used to help inform how to best expand stents as well as to design stents that apply the correct level of strain.

In addition to changing the strain conditions applied, the strain device could also be easily modified to investigate different potential factors in in-stent restenosis. As VSMC have also been implicated in in-stent restenosis[3; 49] those cells could also be investigated for response to stent strut indentation. In addition to cells being tested individually, this model could accommodate full strips of vascular tissue, although they would have to be of a larger species such as porcine, bovine or human. Using strips of tissue would yield a model that could recapitulate how different cell types interact with each other, and where the smooth muscle cells are truly contractile, as opposed to cultured smooth muscle cells that adopt a synthetic phenotype. A model based on full tissue strips would also enable a better simulation of stent strut interaction with blood vessels to understand the role of stenting forces in in-stent restenosis.

### 4.4 Conclusions

MVSC that cannot reorient, and consequently strain avoid, will proliferate when exposed to uniaxial tensile strain. Therefore, MVSC exposed to increased tensile strain could potentially over proliferate causing intimal thickening and even stenosis, thereby suggesting a significant role for MVSC in vascular disease. This research highlights the importance of cell orientation in terms of controlling vascular cell growth. When designing tissue engineered vessels or vascular interventions, such as stenting, which change the mechanical environment in a blood vessel, it is essential to understand not only the strain applied to the underlying substrate, but how this strain translates to the strain sensed by the cells. The strain sensed by cells is dependent upon cellular alignment. Ultimately this alignment-directed sensing of strain could be critical in understanding cell response in vascular disease. By creating a device that allows the application of both static and dynamic stent strut indentation, this paper has demonstrated that MVSC are potentially a dominant cell type in in-stent restenosis.

## 1 Data Availability Statement

The raw data supporting the conclusions of this article will be made available by the authors, without undue reservation.

